# OPA1 mediates cardiac function and metabolism: in silico and in vivo evidence

**DOI:** 10.1101/2024.08.23.605375

**Authors:** Claire Fong-McMaster, Serena M. Pulente, Luke Kennedy, Tyler K.T. Smith, Stephanie Myers, Michel Kanaan, Charbel Karam, Matthew Cope, Ilka Lorenzen-Schmidt, Craig J. Goergen, Morgan D. Fullerton, Miroslava Cuperlovic-Culf, Erin E. Mulvihill, Mary-Ellen Harper

## Abstract

OPA1 is an inner mitochondrial membrane protein that mediates diverse signaling processes. OPA1 is important for cardiac function and protects against cardiac insults such as ischemia reperfusion injury. We sought to further assess OPA1 in human and mouse cardiac pathologies, hypothesizing that OPA1 may also function in a protective manner in chronic heart failure. Bioinformatic analyses of histological and transcript data from the GTEx database indicated that OPA1 expression levels vary in the human heart, where elevated OPA1 transcript levels were correlated with fatty acid, branch chain amino acid and contractile gene signatures. To experimentally assess these correlations, mice with a 1.5-fold whole body OPA1 overexpression (OPA1-OE) were subjected to transverse aortic constriction surgery and displayed improved 2D and 4D cardiac functional parameters compared to WT mice. OPA1-OE mice had no induction of fibrotic transcript markers and displayed sustained transcript levels of fatty acid, branch chain amino acid and contractile markers. Maximal oxidative capacity was sustained in both WT and OPA1-OE cardiac myofibers post-TAC. These results further demonstrate the important role of OPA1 in mediating cardiac function and highlight protective signaling pathways.

## Introduction

Heart failure affects over 50 million people worldwide with increasing rates attributable to complex etiologies and risk factors^1,2^. Many pharmacological approaches exist for disease management which include the use of ACE inhibitors, beta blockers, diuretics and SGLT2 inhibitors^3^. These therapies combined with nonpharmacological approaches such as diet and exercise require complex individualized care that is further complicated by comorbidities. Given this landscape, characterizing the underlying mechanisms which contribute to reduced cardiac function at each disease stage is critical to designing new and complementary therapeutic approaches. Emerging evidence points to the potential role of mitochondrial-targeted therapeutics to treat heart failure^4,5^. Changes in mitochondrial metabolism are evident throughout the progression of heart failure and largely include irregular substrate metabolism, alterations in mitochondrial energetics, reactive oxygen species overload and an imbalance in mitochondrial dynamics^6^.

Mitochondria make up 30% of cardiomyocyte volume and undergo cycles of fusion and fission in response to altered energy status or cellular stressors^7^. This dynamic process involves inner mitochondrial membrane (IMM) and outer mitochondrial membrane (OMM) proteins, as well as proteins that tether to adaptor proteins on the OMM^8^. Fission is largely driven by this latter class of proteins, in which dynamin-related protein 1 is recruited to the OMM and constricts mitochondria leading to mitochondrial fission. Fusion is driven by three GTPases, mitofusin 1 and 2 on the OMM, as well as optic atrophy protein-1 (OPA1) on the IMM. In addition to mediating mitochondria fusion, OPA1 is linked to oxidative phosphorylation efficiency, cristae morphology, apoptotic signaling, and calcium signaling^9,10^.

In both human and pre-clinical models of heart failure, OPA1 protein abundance has been shown to be lower relative to healthy controls^11,12^. OPA1 heterozygous mice exhibit impairments in contraction, cardiac output and contractile response when aged or subjected to pressure-overload^13,14^. Underlying these functional changes are differential calcium handling within cardiomyocytes, abnormal mitochondrial ultrastructure, and altered electron transport chain activity. Conversely, increased OPA1 protein levels have been shown to be protective during acute myocardial injury by maintaining cristae structure and minimizing apoptotic signaling^15^. Beyond protein abundance, OPA1 processing by the proteases YME1L and OMA1 impacts cardiac function^16–18^. Mitochondrial substrate utilization is closely linked to mitochondrial dynamics with evidence highlighting the role of fatty acids in increasing YME1L expression, thereby limiting OPA1 cleavage and mitochondria fragmentation^19^. OPA1 is also sensitive to redox modifications in the myocardium which can contribute to mitochondrial dysfunction^20^. Despite knowing that decreased OPA1 worsens cardiac function, it is unknown how a subtle increase in expression, and importantly one which could be therapeutically achievable, would influence cardiac function in a chronic heart failure state.

Given the evidence implicating OPA1-mediated mechanisms in various heart failure models, we sought to assess the molecular mechanisms that link OPA1 and cardiac pathologies. Using human and mouse *in vivo* data, we confirm the role of OPA1 in mediating cardiac function with links to fatty acid, branch chain amino acid and contractile pathways.

## Results

### OPA1 transcript level correlates with metabolic, contractile, and hypertrophic pathways in human left ventricle tissue

OPA1 expression levels are decreased in human heart failure^11^, but a correlation of OPA1 expression and histopathology features of cardiac pathologies has not been characterized. To investigate this, we first assessed human gene expression profiles using the Genotype-Tissue Expression (GTEx) Portal^21,22^ (Fig 1A). RNA-seq data from GTEx left ventricle (LV) samples were filtered based on *OPA1* expression. Top and bottom quintiles of *OPA1* expression were defined from all samples under the same cause of death category (ventilator case, n=246 samples). Interestingly, a range of *OPA1* transcript expression was observed (Fig 1B), with a median log_2_fold change difference of 1.72 between top and bottom quintile and only small differences in overall gene expression values between the groups (top and bottom median normalized transcript per million (TPM) values: 0.01338 and 0.01344, respectively. p-value=0.000109) (Fig S1A).

**Figure 1.**
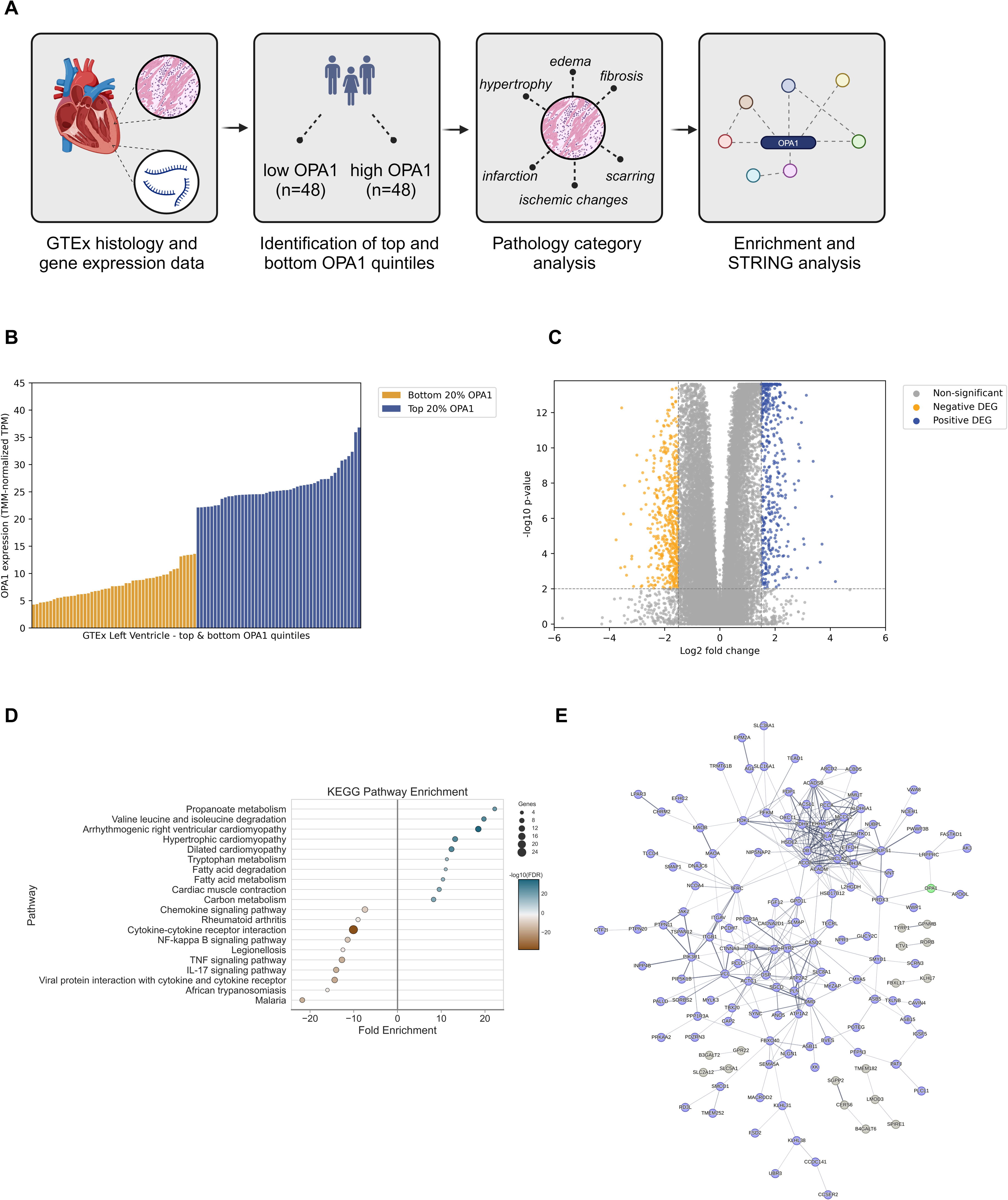
OPA1 expression in top and bottom quintiles of GTEx left ventricle samples. (A) Schematic of GTEx data processing and analysis. **(B)** Normalized expression values of OPA1. **(C)** Differential gene expression between top and bottom quintiles of OPA1 expression. Significance thresholds are |1.5| median log_2_ fold change and 0.01 adjusted p-values. **(D)** Enriched KEGG pathways in positive (blue) and negative (orange) DEGs between top and bottom expressors. **(E)** Interaction network of positive DEGs with OPA1 (green). Central cluster nodes are coloured blue, non-central clusters are grey. Edges are defined by interaction confidence through STRING evidence sources.

To explore the relationship between *OPA1* transcript expression and cardiac pathologies, we analyzed histological data from the GTEx portal. Relevant pathology categories in the LV histology samples such as fibrosis, as identified by pathologists^23^, were selected and their rates of occurrence compared between the top and bottom *OPA1* expressors, as well as between all samples (Table 1). Fibrosis was identified more frequently in the lowest *OPA1* expressing samples compared to higher expressing *OPA1* LV samples (p=0.0049), and all LV samples (p=0.0203), though fibrosis was not significantly lower in top samples compared to all samples (p=0.1784).

**Table 1.** Incidence of pathology categories in top and bottom OPA1 quintiles of GTEx left ventricle samples.

|  | Pathology category |  |  |  |  |  |
| --- | --- | --- | --- | --- | --- | --- |
|  | fibrosis | hypertrophy | infarction | ischemic_changes | edema | scarring |
| Bottom OPA1 | 23 | 2 | 2 | 13 | 0 | 1 |
| Top OPA1 | 9 | 4 | 2 | 16 | 0 | 1 |
| Overall | 72 | 12 | 7 | 62 | 1 | 5 |
| Bottom proportion | 0.4792 | 0.0417 | 0.0417 | 0.2708 | 0 | 0.0208 |
| Top proportion | 0.1875 | 0.0833 | 0.0417 | 0.3333 | 0 | 0.0208 |
| Overall proportion | 0.2939 | 0.0490 | 0.0286 | 0.2531 | 0.0041 | 0.0204 |

Given the role of mitochondrial metabolism and dynamics in cardiac aging^24,25^, we assessed any potential differences in *OPA1* transcript across all ages. We observed a trend of elevated *OPA1* expression in LV samples aged 20-49 compared to LV samples isolated from those aged 50-79 (p=0.0639 in low quintile and p=0.0479 in top quintile) (Fig S1B).

To explore the underlying molecular signatures of LV in the top and bottom *OPA1* quintiles, we identified gene expression changes associated with changes in *OPA1*. Three approaches including differential expression analysis, partial least squares discriminant analysis (PLS-DA)^26^ and Relief-based analysis were used to identify statistically relevant genes^27,28^. Using differential expression analysis, 441 and 488 genes were positively and negatively differentially expressed between the *OPA1* quintiles in LV (Fig 1C). Two machine learning methods (PLS-DA and TuRF algorithms) identified 3582 and 3427 important feature genes, respectively, using variable importance projection (VIP) scores^29^ for PLS-DA (Fig S1C) and relevance thresholds for TuRF. In comparing *OPA1* high to low expression sample groups, 229 positive differentially expressed genes (DEGs) were identified by all three methods, and only 31 negative DEGs were found by all methods; indicating a more consistent association with increased *OPA1* expression compared to decreased expression.

Gene set enrichment was applied to these 229 genes from the conserved gene sets, for KEGG (Fig 1D) and GO Cellular Component terms (Fig S1D). Multiple metabolic pathways were associated with *OPA1* expression including propanoate, branched-chain amino acid, and fatty acid metabolism. Additionally, an enrichment in terms for cardiac muscle contraction and adrenergic signaling was clearly associated with increased *OPA1* expression. Enrichment analysis for the 31 negative conserved DEGs could not be determined due to the limited gene set size. Enrichment in the 488 statistically down-regulated genes identified by differential expression analysis were reported (Fig 1D). We identified immune responses amongst the negatively enriched terms. Amongst enriched cellular component GO terms (Fig S1D), the sarcoplasmic reticulum, TCA complex, and contractile regions (Z disk, I band) are associated with *OPA1* while hemoglobin is amongst the negatively associated terms.

STRING^30,31^ interaction networks identified relationships between significant genes through various associations, including protein-protein interaction, text mining, and homology. For the 229 upregulated DEGs identified by our differential expression analysis and machine-learning methods, 144 genes have interactions and 126 of these are connected through a central cluster (Fig 1E). *OPA1* is connected to this central cluster via the *LRPPRC, PRDX3*, and *APOOL* genes. These data provide evidence for potential protein-protein interactions between *OPA1* and many of the proteins within the enriched functional pathways.

### OPA1 overexpression maintains cardiac functional parameters compared to wildtype mice in response to pressure overload

Given the correlation between human *OPA1* mRNA expression and known metabolic pathways involved in heart failure progression, we followed up with *in vivo* work to assess how increased levels of OPA1 would impact cardiac function. To do so, we employed OPA1 overexpressing (OPA1-OE) mice, which exhibit a 1.5-fold whole body overexpression^15^, and subjected them to a transverse aortic surgery^32^ (TAC), where banding is placed around the descending aorta designed to more closely mimic human pathology. Mice subjected to TAC surgery exhibited a heart failure phenotype at 12 weeks post-TAC when compared to sham-operated mice, with statistically significant increases in LV mass, LV end diastolic diameter (LVEDD), and LV end systolic diameter (LVESD) but no changes in lung weight or lung water content (Fig S2A-E).

Mice of both genotypes were randomly assigned to receive TAC surgery or sham surgery (Fig 2A). At 5-, 8- and 12-weeks post-TAC, OPA1-OE mice had higher ejection fraction evaluated by long axis B mode echocardiography compared to WT mice, indicative of improved cardiac remodeling in response to pressure overload (Fig 2B-C). This was also supported by short axis imaging (Fig 2D) which also showed higher ejection fraction and cardiac output in OPA1-OE TAC mice compared to WT TAC mice (Fig S3A-B). Cardiac structural remodeling was evident in both WT and OPA1-OE TAC mice as both genotypes had significant increases in the heart weight/tibia length (HW/TL) and heart weight/body weight (HW/BW) ratios compared to sham mice (Fig 2E). While both WT and OPA1-OE groups exhibited modest increases in EDV with TAC, OPA1-OE EDV increased to a lesser extent (Fig 2F). The same trend held true for ESV (Fig 2G). Conversely, there were no genotypic changes observed in stroke volume or cardiac output (Fig 2H-I). These data suggest that increased OPA1 expression during pressure overload induced heart failure in mice is protective in maintaining cardiac function.

**Figure 2.**
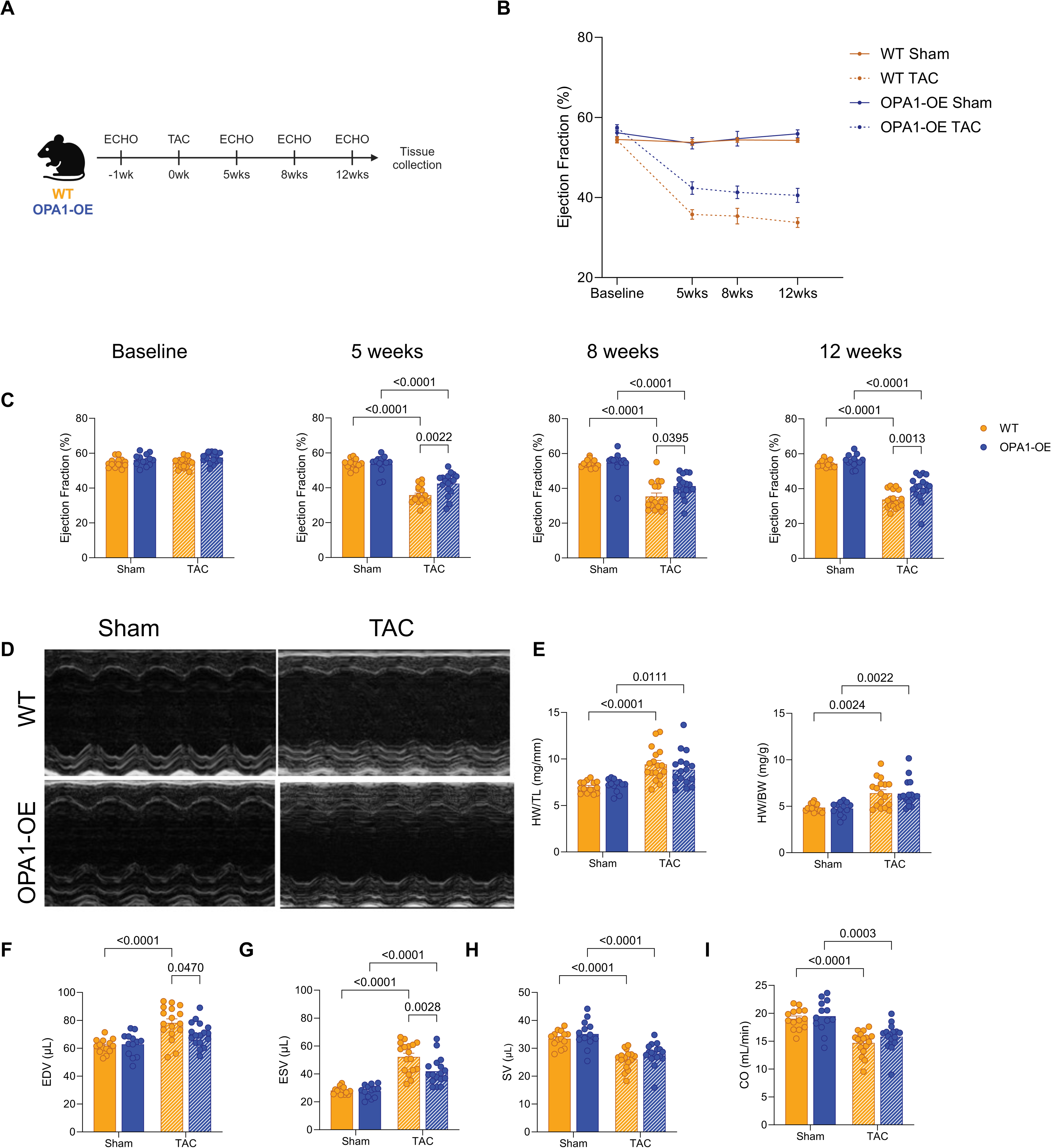
**OPA1 overexpression mediates cardiac functional protection in a pressure-overload model**. **(A)** Schematic of experimental design and study imaging timepoints. **(B)** Ejection fraction measured by 2D parasternal long axis overall timepoints. **(C)** Ejection fraction measured by 2D parasternal long axis at baseline, 5 weeks, 8 weeks, and 12 weeks post-TAC. **(D)** Representative short axis M-mode images. **(E)** Heart weight to tibia length ratio (HW/TL) and heart weight to body weight (HW/BW) ratio at 12 weeks post-TAC. **(F-I)** 2D parasternal long axis measurements including end diastolic volume (F), end systolic volume (G), stroke volume (H) and cardiac output (I). All data are represented as the mean ± SEM. P values calculated using a two-way ANOVA with Tukey’s post-hoc analysis for (C,E-I).

### 4D strain analysis identifies differences in global myocardial strain in response to pressure overload

While 2D imaging can detail cardiac remodeling, volumetric measurements are estimated from only one plane of view. To fully characterize volumetric changes in response to TAC surgery, we employed 4D echocardiography, which collects a 3D image at every time point along the short axis using a step motor, allowing for analysis of myocardial kinetics and motion. Using a custom strain technique^33,34^, we assessed global strain parameters at 12 weeks post-TAC (Fig 3A). Global circumferential strain was decreased in both WT and OPA1-OE mice compared to their respective sham mice (Fig 3B). There was a trend for decreased longitudinal strain in WT TAC mice compared to WT sham mice and no significant changes between other groups (Fig 3C). Surface area strain was increased in both WT and OPA1-OE TAC mice, but OPA1-OE TAC mice maintained lower strain compared to WT TAC mice (Fig 3D). Finally, transmural (or radial) strain was decreased in WT TAC mice compared to WT sham mice with a trend for decreased transmural strain in OPA1-OE TAC mice compared to OPA1-OE sham mice (Fig 3E).

**Figure 3.**
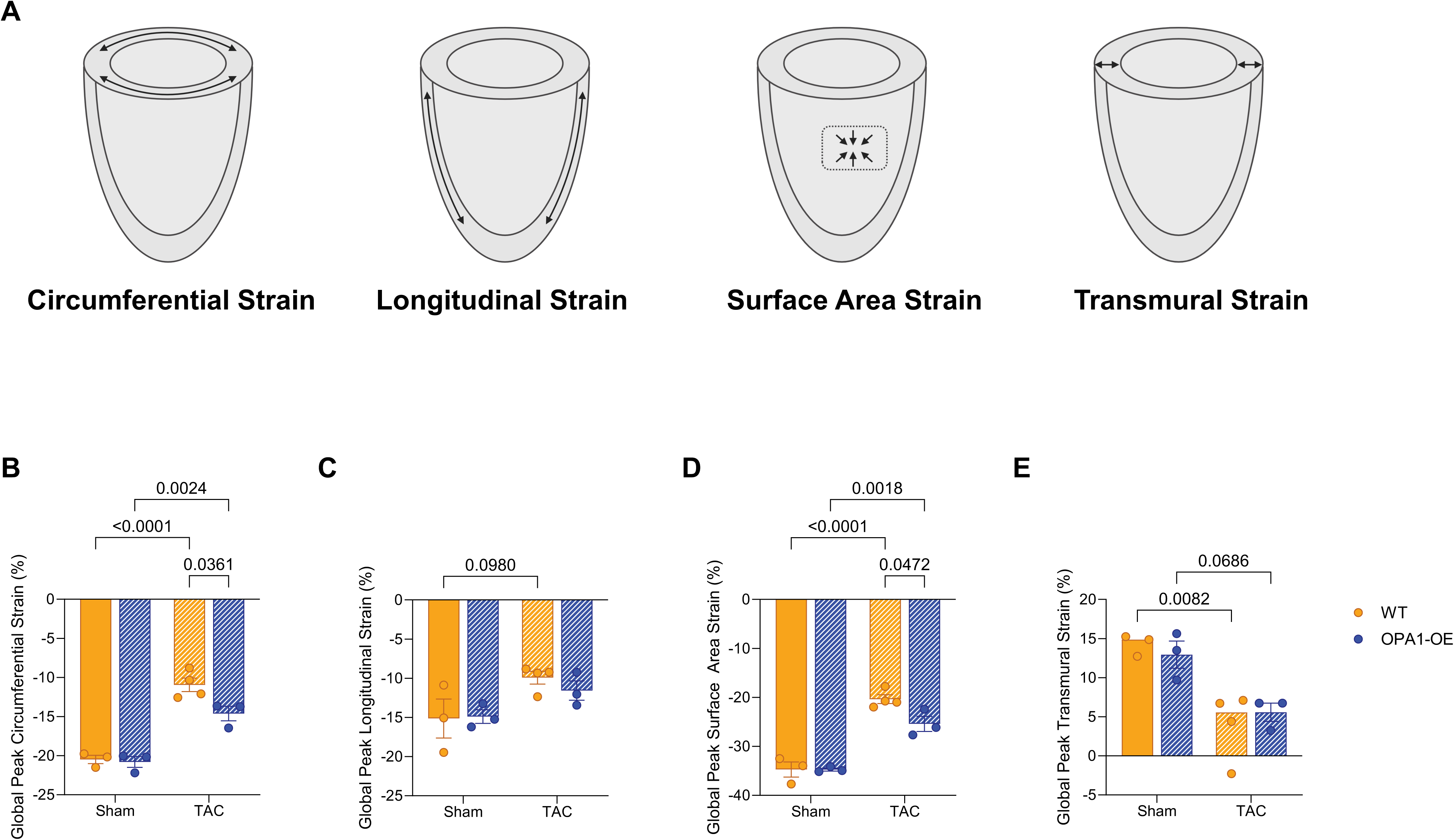
4DUS strain analysis identifies changes in global strain parameters post-TAC in WT and OPA1-OE mice. (A) Schematic of global strain orientations. **(B-E)** Global strain parameters including circumferential (B), longitudinal (C), surface area (D), and transmural (E). All data are represented as the mean ± SEM. P values calculated using a two-way ANOVA with Tukey’s post-hoc analysis for (C-F).

### TAC induces trends for hypertrophic and fibrotic remodeling in WT mice

To address what could mediate the cardioprotective effects of OPA1 overexpression, we assessed markers of hypertrophic and fibrotic remodeling at 12 weeks post-TAC. Cardiac fibrosis is central to the transition from hypertrophy to decompensated heart failure. In classical, more aggressive TAC models, extensive fibrosis is measurable as early as 2 weeks^35,36^. However, despite having significant decreases in cardiac function, we observed no changes in interstitial and perivascular fibrosis in our more physiological TAC model (Fig 4A-C). In line with our minimal fibrotic phenotype, there were only trends for increased levels of hypertrophy markers including *Nppb* in WT TAC mice (Fig 4D). Similarly, TAC induced modest increases in fibrosis markers including *Tgfb1, Col1a2,* and *Ctgf* in WT mice compared to sham mice (Fig 4E). However, this was less apparent in OPA1-OE mice.

**Figure 4.**
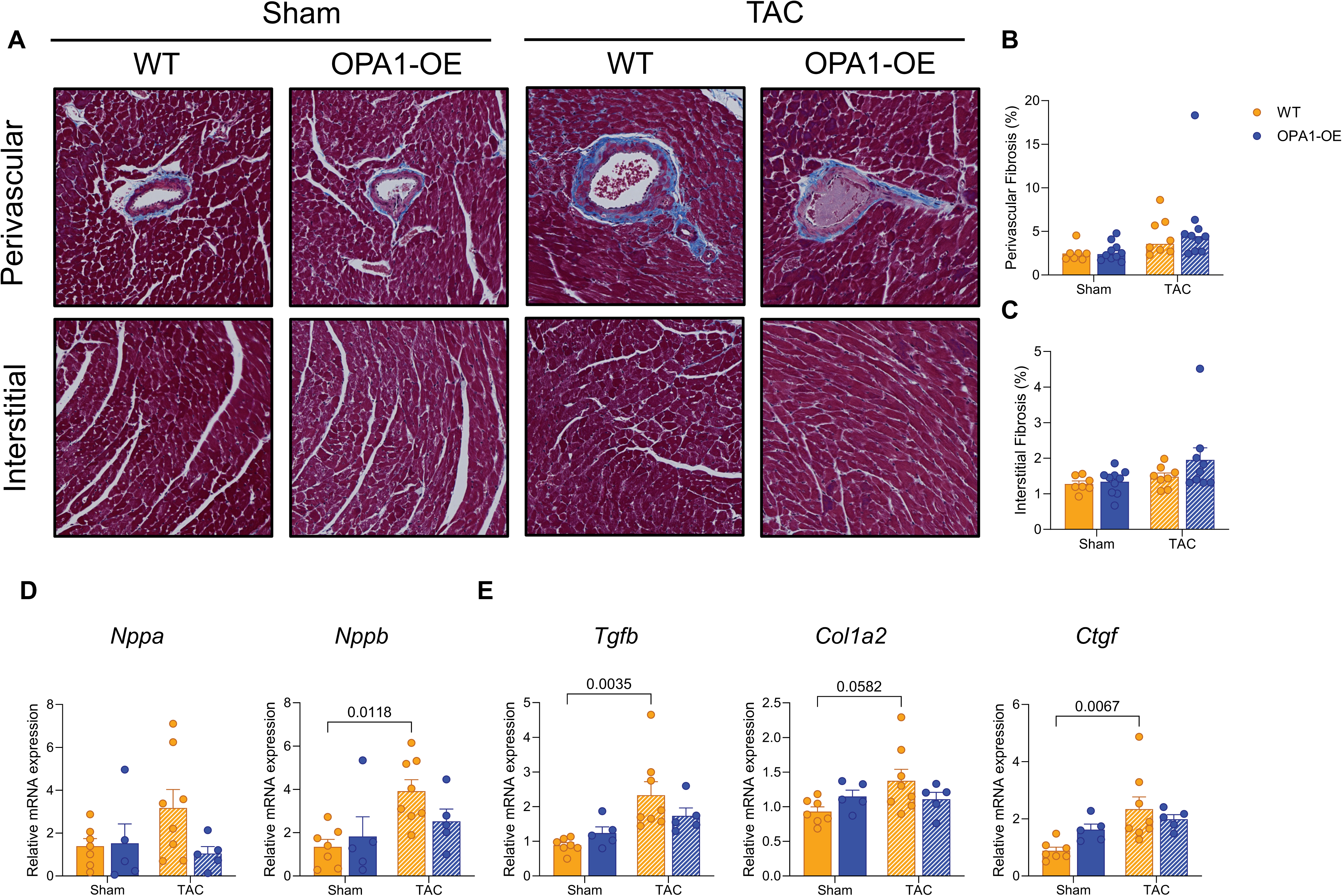
Hypertrophic and fibrotic genes are upregulated in WT but not OPA1-OE TAC mice. (A) Representative images of trichrome staining of LV cross sections. **(B-C)** Quantification of perivascular (B) and interstitial fibrosis (C) from trichrome-stained slides. **(D-E)** Relative mRNA expression of hypertrophy markers (*Nppa* and *Nppb*) and fibrosis markers (*Tgfb*, *Col1a2*, and *Ctgf*) in cardiac tissue at 12 weeks post-TAC. All transcript data are expressed as relative to the geometric mean of *B2m* and *Eef1e1.* All data are represented as the mean ± SEM. P values calculated using a two-way ANOVA with Tukey’s post-hoc analysis for (B-E).

### Transcriptional changes between OPA1-OE and WT TAC mice

Metabolic remodeling through the progression from hypertrophy to heart failure is complex and involves changes in substrate uptake, utilization and oxidative metabolism^37^. To see if these metabolic pathways were relevant in our model, we assessed transcript levels of fatty acid oxidation and BCAA metabolism markers. Markers of fatty acid metabolism including *Cd36* and *Acadm* were modestly decreased in WT TAC mice compared to WT sham mice, and *Cd36* was significantly lower in WT TAC compared to OPA1-OE TAC mice (Fig 5A-C). BCAA markers including *Bcat2* and *Bckda* were lower in WT TAC mice, but *Bcat2* remained elevated in OPA1-OE TAC mice (Fig 5D-E). Another pathway that was identified to be correlated with OPA1 expression was cardiac contraction. Given this, we measured *Atp2a2* transcript levels, which were significantly higher in OPA1-OE TAC mice compared to WT TAC mice (Fig 5F).

**Figure 5.**
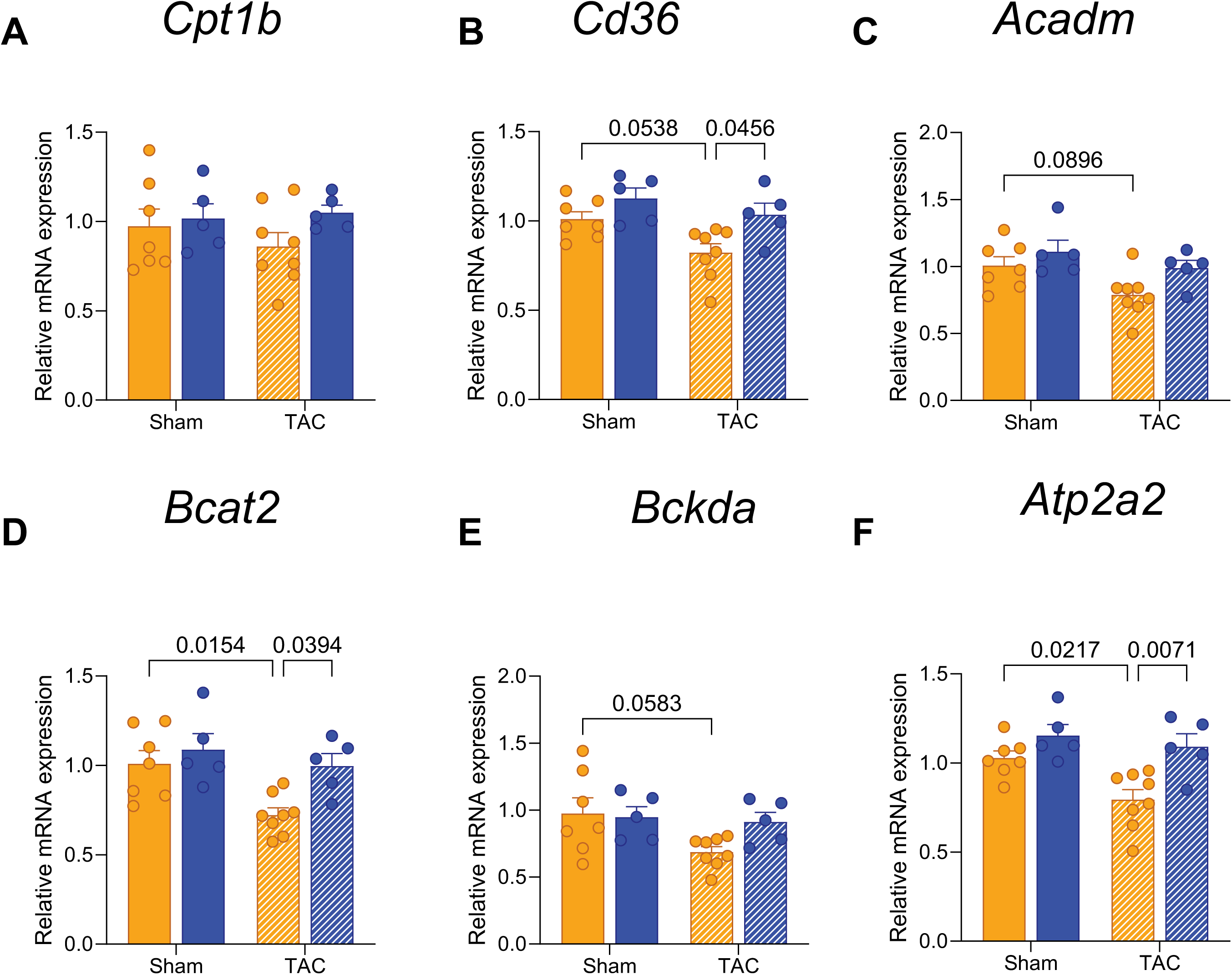
Expression of metabolic and contractile-related genes are higher in OPA1-OE TAC mice. (A-F) Relative mRNA expression of fatty acid metabolism (*Cpt1b, Cd36*, and *Acadm*) (A-C), branch chain amino acid metabolism (*Bcat2* and *Bckda*) (D-E) and contraction (*Atp2a2*) (F) in cardiac tissue at 12 weeks post-TAC. All transcript data are expressed as relative to the geometric mean of *B2m* and *Eef1e1.* All data are represented as the mean ± SEM. P values calculated using a two-way ANOVA with Tukey’s post-hoc analysis for (A-F).

### Sustained oxidative capacity at 12 weeks post-TAC

We next wanted to assess the implications of OPA1 overexpression to mediate changes in oxidative capacity of the heart. Analyses of maximal oxygen consumption in permeabilized LV myofibers were measured in response to saturating concentrations of palmitoyl carnitine/malate, glutamate/pyruvate, and succinate. Surprisingly, there were no changes between TAC and sham mice of either genotype (Fig 6A-E). Simultaneous measurement of hydrogen peroxide emission to assess reactive oxygen species, indicated no changes between TAC or sham mice of either genotype (Fig 6F-J). In addition, we measured mitochondrial DNA/nuclear DNA ratios and citrate synthase activity as indicators of mitochondrial content and observed no differences (Fig 6K-M).

**Figure 6.**
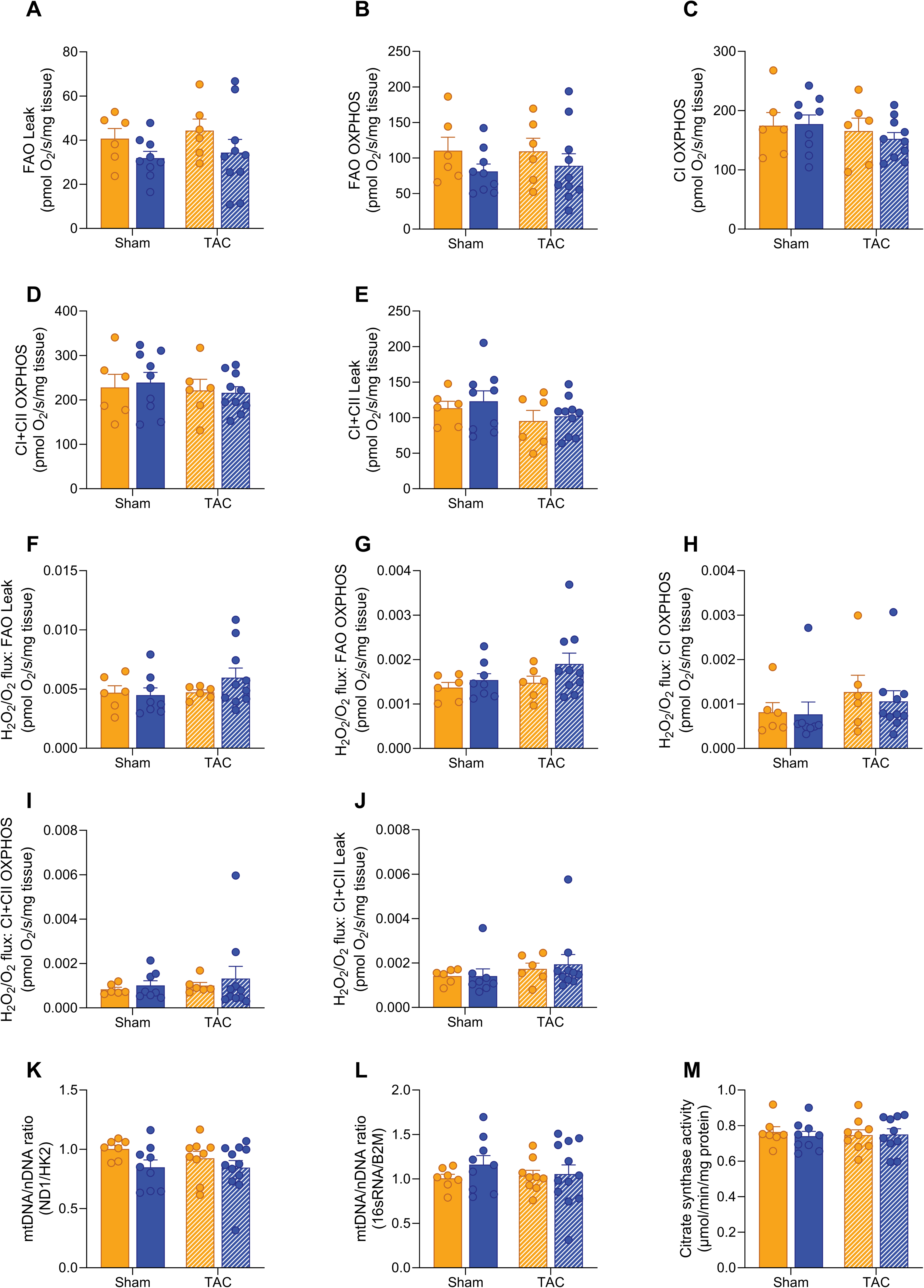
Mitochondria maximal oxidative capacity and hydrogen peroxide emission is unchanged in permeabilized myofibers post-TAC. (A-E). High resolution respirometry measures of oxygen consumption in permeabilized cardiac myofibers. **(F-J)** Hydrogen peroxide emission normalized to oxygen flux. (M-N) Ratio of mitochondrial DNA to nuclear DNA measured using primers for *Nd1* and *Hk2* (K) and 16sRNA and B2M (L). (M) Citrate synthase activity measured in protein homogenate of cardiac tissue. All data are represented as the mean ± SEM. P values calculated using a two-way ANOVA with Tukey’s post-hoc analysis for (A-F).

## Discussion

In our work, we present the first evidence for the correlation between OPA1 expression levels and advanced characteristics of cardiac pathology in humans. Our analyses of publicly available transcriptomics data from the GTEX consortium indicates that OPA1 mRNA levels vary significantly in left ventricular tissue in the human heart. Higher OPA1 expression was correlated with elevated fatty acid and branched chain amino acid metabolism, and cardiac contractile pathways. We investigated this observation using a mouse model of modest OPA1 overexpression, where we observed higher cardiac function and kinetic motion in response to pressure-overload surgery, sustained transcript levels of fatty acid, BCAA and contractile genes with no changes in maximal oxidative capacity.

Heart failure is a chronic disease with recent efforts elucidating transcriptional signatures representative of stages of disease progression^38^. Our transcriptomic analysis of OPA1 levels in the GTEx database identify increased fibrosis in human LV samples within the lowest OPA1 expression quintile. These findings highlight a correlation between OPA1 and one of the central pathologies in heart failure. This is in line with the incidence of cardiac pathologies in patients with autosomal dominant optic atrophy, a disease where up to 70-90% patients present with a mutation in OPA1^39,40^. These clinical reports have noted early-onset myocardial infarction^41^, arrhythmia^42,43^, and hypertrophic cardiomyopathy^42^, calling for the recommendation for further investigation into the connection between OPA1 mutations and cardiac events^42^.

Several methodologies are employed to induce heart failure in mice. Most TAC studies use transverse aortic banding with a 27-gauge needle, which induces rapid decreases in cardiac function and subsequent fibrotic remodeling evident within 2 weeks. Previous literature suggests that the placement of the aortic constriction will affect the degree of pressure overload^32^ and that the degree of constriction affects the severity of hypertrophic and fibrotic remodeling^36^. We used a less severe model of heart failure, in which the descending aorta is constricted using a 26-gauge needle. We opted to use the descending aortic banding with a 26-gauge needle^44^ to investigate chronic changes and minimize animal mortality. This explains our minimal fibrotic phenotype in both TAC groups and lack of progression to decompensated heart failure.

While the link between cardiac function and OPA1 has been assessed using heterozygous models^13,14,45^, an OPA1 overexpression model^15^, and models modulating upstream proteases or cleavage sites^16–18^, our work is the first to increase OPA1 levels during a chronic *in vivo* heart failure model. Mild OPA1 overexpression maintained cardiac functional parameters including ejection fraction and fractional shortening, as well as longitudinal and circumferential strain post-TAC compared to WT mice. These results align with data showing that OPA1 heterozygous mice are more sensitive to pressure overload and ischemia/reperfusion^13,45^. Original characterization of the OPA1-OE mouse model^15^ identified no baseline changes in cardiac function or structure at 5 months and protection from ischemia^15^. When aged to 9 months, these mice develop non-pathological hypertrophy as indicated by an increased HW/BW ratio. These results may explain how the OPA1-OE mice subjected to TAC resulted in increased HW/BW and HW/TL ratios yet higher functional parameters compared to WT TAC mice. OPA1 has also been implicated in protective cardiac remodeling in response to various treatment interventions *in vivo* such as empagliflozin^46,47^, irisin^48^ and paeno1^49^. This and other studies highlight OPA1 as a key mediator of protective cardiac phenotypes likely due to several mechanisms including its established role in mediating cristae remodeling and apoptotic signaling, mitochondrial fusion, mitochondrial supercomplexes, oxidative metabolism, and calcium homeostasis.

Metabolic changes in heart failure involve alterations in substrate uptake and utilization. Our GTEx analysis and subsequent mRNA transcript data identified that OPA1 is important for regulating fatty acid metabolism. These data further support the link between mitochondrial fusion and fatty acid oxidation previously identified in cardiac tissue^19^. Branch chain amino acid catabolism is another central metabolic pathway known to be decreased in heart failure and pressure-overload mouse models^50,51^. We identified sustained transcript levels of certain BCAA enzymes in OPA1-OE TAC mice compared to WT TAC mice indicating a protective metabolic phenotype within our model. Another indicator of cardioprotection in the OPA1-OE TAC model was increased *Atp2a2* transcript levels compared to WT mice. Calcium signaling is critical for maintaining the function of contractile machinery in the heart^52^. Defects in calcium kinetics have been observed in Opa1 heterozygous mice^13,14^. These data further support the importance of OPA1 in oxidative metabolism and calcium signaling in the heart.

Heart failure is typically characterized by an “energy deficit” with decreases in electron transport chain activity or maximal oxidative capacity. Given this, we assessed various bioenergetic pathways, identifying no changes in oxidative capacity between sham and TAC mice in response to the sequential addition of fatty acid, complex I, and complex II substrates. In addition, we did not identify changes in mitochondrial content. Previous *in vivo* work has shown that TAC induces changes in maximal oxidative capacity, mitochondrial enzymatic activities and the expression of mRNA and protein levels of mitochondrial DNA^53,54^. Although these changes may be largely driven by decreases in overall mitochondrial content, our conflicting experimental results highlight important methodological considerations in assessing bioenergetic function. To assess maximal respiratory capacity, we used the sequential addition of saturating concentrations of various substrates. However, this does not fully represent *in vivo* steady state conditions, which may have been altered post-TAC. The subpopulations of mitochondria within cardiac tissue including the subsarcolemmal and interfibrillar mitochondria are differentially affected by pressure-overload^55^ with interfibrillar mitochondria showing more significant decreases in respiratory capacity. In our experiments, we did not separate out these two populations so any functional differences may be masked by measuring the total myofiber response to the substrates. In addition, assessing myofibers bioenergetic capacity instead of isolated mitochondria may explain the lack of functional deficits that we observed^56^. In human work, data has suggested that oxidative capacity is reduced in *ex* vivo myofiber preparations from ventricular tissue of patients with heart failure^57^. These data were supported by decreases in mitochondrial content measures, which may explain the functional changes. Conflicting work has identified preserved respiratory function in isolated mitochondria from human LV tissue^58^ highlighting the disease heterogeneity and the important methodological considerations when assessing bioenergetic function. Our *in vivo* TAC model therefore likely represents a less severe form of heart failure that occurs before significant changes to total mitochondrial pools.

In conclusion, this study highlights the importance of OPA1 in mediating cardioprotective mechanisms. Our bioinformatic approaches using publicly available human data identified correlations between low *OPA1* expression and cardiac tissue fibrosis as well as the link between *OPA1* and metabolic pathways. Furthermore, *in vivo* in mice support a role for OPA1 in mediating metabolic and contractile signaling. Overall, we provide further evidence for the importance of OPA1 in cardiac function and elucidate gene signatures that underlie this protection.

### Limitations

Our study has various limitations which need to be addressed. First, all experiments were performed in male mice. As such we cannot conclude the applicability of these findings in female mice. Secondly, aortic banding around the descending aorta results in difficult imaging of the pressure gradient across the constriction site. Careful use of ultrasound probes with longer scan depth (eg. MX250S) is required to obtain resolution of the descending aortic arch. In addition, the use of other non-invasive measures for surgical validation such as ECG measures will be critical in the future use of this model. The OPA1-OE mouse model exhibits a whole-body overexpression of OPA1, so we cannot conclude that cardiac OPA1 is solely responsible for the protective phenotype observed post-TAC. In addition, we did not assess OPA1 processing and isoforms, which has been elucidated in other cardiac studies^16–18^. All endpoint experiments in our study were conducted at 12 weeks post-TAC, which may have limited our findings as mitochondrial content have been shown to transiently increase post-TAC^59^. Instead, we focused on an extended timepoint at 12 weeks post-TAC to assess the role of OPA1 in a more chronic disease state.

## Methods

### Bioinformatic analyses

All bioinformatic analyses were carried out using Python (Version 3.8.18) with figures and plots generated via the matplotlib^60^ and seaborn^61^ libraries. All statistical tests were performed through their respective Scipy method^62^. All data and code for these analyses are available in our Github repository: https://github.com/lkenn012/OPA1_heart_GTEx.

### GTEx gene expression analysis

LV GTEx RNA-seq data were collected (TPM, v8) via the GTEx Portal API to identify samples within top and bottom quintiles of OPA1 expression. Samples without expression data for OPA1 were first removed then TPM values were normalized by trimmed mean of M-values (TMM)^63^ using the conorm library (https://pypi.org/project/conorm/). Samples were selected irrespective of age with an equal number of males and females selected. Initially, all RNA-seq samples were considered (n=433), though after selecting top and bottom quintiles by OPA1 expression it was observed that top quintile consisted exclusively of individuals classified as “Ventilator cases” within the Hardy scale cause of death categorizations^64^. Though this represents roughly half of all GTEx individuals, to avoid possible confounding factors in the analysis quintiles were again selected from just this subset of samples (n=246). From this subgroup of GTEx samples, top and bottom quintiles were selected by OPA1 expression (n=48 per group).

### Histology images pathology

LV histology images for OPA1 quintiles were obtained from the National Cancer Institute’s Biospecimen Research Database^65^. Histology data were downloaded directly from the GTEx histology viewer (https://gtexportal.org/home/histologyPage) for comparison of pathologies noted in OPA1 quintiles. Occurrences of pathology categories of interest in each group were compared by chi-squared tests.

### Differential gene expression analysis

Differentially expressed genes between OPA1 quintiles were determined as those genes with TMM-normalized expression values with absolute median log_2_fold change values greater than 1.5 and false discovery rates < 0.01 as computed by Wilcoxon’s rank sum test after adjusting for multiple testing by Benjamini-Hockeberg correction (Statsmodels^66^). Genes with unusually large log_2_fold changes (>8 or <-8) were also excluded as these likely come from comparisons to samples with no expression of the gene.

### Supervised machine learning feature selection

Relevant genes for OPA1 expression were determined by PLS-DA^26^ for classifying top and bottom quintile GTEx expressors, or as predictors of OPA1 expression across top and bottom quintiles by the multi-round Relief-based approach, TuRF^27,28^. The PLS-DA algorithm was implemented using the scikit-learn^67^ “PLSRegression” function, with default parameters using z-score normalized gene expression values as features and quintile groups as targets. The multi-round TuRF algorithm^28^ was implemented via skrebate^68^ with default parameters (using all samples as neighbours for computing weights) and the ReliefF algorithm^69^ as a base (single-round) estimator to predict OPA1 expression across top and bottom expressors.

Relevant features were selected according to peak model performance over VIP score thresholds^29^ for PLS-DA and according to relevance threshold for Relief features.

### Gene-set enrichment analysis

Enriched functional terms in important genes for OPA1 expression, defined as the overlapping gene set identified by differential expression and feature selection methods, were identified by ShinyGO^70^ (version 0.80). All genes expressed in GTEx LV samples were used as background and an FDR cutoff of 0.01, and minimum pathway size of 10 were applied to KEGG^71^ and gene ontology^72^ term enrichment analysis to identify the top 10 enriched terms and corresponding values for each.

### Interaction networks

The STRING database^30,31^ was used to construct interaction networks between the overlapping gene sets in our GTEx analysis. Networks were constructed based on a medium confidence threshold defined by STRING using all interactions sources, with edge thickness proportional to the level of interaction evidence from the various sources. Nodes are coloured by clusters and OPA1 is highlighted.

### Animal studies

All experiments involving mice were conducted in accordance with the principles of the Canadian Council of Animal Care and were approved by the Animal Care Committee of the University of Ottawa (UOHI-AUP-2909). Male and female OPA1-OE mice were a kind gift from Dr. Luca Scorrano^15,73^. Mice were genotyped using the following primers:

Transgenic allele (expected size 1200bp):

M_OPA1TG_FWD: 5′-GCA ATG ACG TGG TCC TGT TTTG-3′

M_OPA1TG_REV: 5′-GAT AGG TCA GGT AAG CAA GCA AC-3′

Wildtype allele (expected size 400bp):

M_WT_FWD: 5′-CTC CGG AAA GCA GTG AGG TAA G-3′

M_WT_REV: 5′-GAG GGA GAA AAA TGC GGA GTG-3′

Animals were housed in 12hr/12hr light cycle with access to standard chow and water.

### Transverse aortic constriction model

Pressure-overload was induced using a modified transverse aortic constriction model. First, the mice were prepared with three analgesics – buprenorphine (0.05mg/kg S.C. 60 minutes prior to surgery; Meloxicam, 1 mg/kg, S.C. 60 minutes prior to surgery and daily for 3 consecutive days; Buprenorphine Slow Release (lasting 72 hours), 1.2 mg/kg, S.C. prior to start of surgery). 2-4% Isoflurane was used as the anesthetic throughout the surgical operation. Mice were intubated using a 20-gauge intravenous catheter attached to a ventilator set to 130-150 breaths/min with a tidal volume of 0.2mL. Once a sterile incision site was prepared, a transverse incision was made and both layers of the thoracic muscles were separated. The chest cavity was opened at the level of the second intercostal space and a chest retractor was used to spread the opening. The descending aorta was bluntly dissected out visually and a 6-0 silk was passed around the aorta using a Surgipro needle. The silk was tightened around a blunted 26g needle placed parallel to the aorta and secured with two knots. The chest retractor was removed, and the chest cavity was closed with a 6-0 suture tie. Mice remained on the ventilator until spontaneous breathing and received 0.5 mL 37°C S.C. saline injection. Mice were monitored in an incubator with supplemental oxygen (1-2 L/min) at 30°C and then returned to the standard housing rooms after recovery.

### 2D and 4D Echocardiography

Cardiac function was measured using a Vevo3100 System with an MX400 transducer (FUJIFILM VisualSonics, Toronto). Mice were anesthetized using 1.5 −3% isoflurane and 1.5 L/min oxygen. Heart rate, respiratory rate and temperature were monitored throughout all 2- and 4-dimensional imaging. 2-dimensional images in PLAX B mode and SAX B and M mode were taken at baseline, 5 weeks, 8 weeks and 12weeks post-TAC surgery. Vevo LAB (5.8.2) was used for all 2D analyses.

4D mode was used to acquire SAX 4D images using a step size of 0.127 mm and frame rate of 400 Hz. 4D strain parameters were analyzed as previous described^33,34,74^ using a custom MATLAB (MathWorks, USA) graphical user interface.

### Histological analyses

Heart tissue was fixed in 10% formalin and processed for paraffin-embedding. 4 µm sections were stained with Masson’s trichrome to assess collagen. ImageJ Software and the Color Deconvolution tool was used to quantify interstitial and perivascular fibrosis.

### High resolution respirometry in permeabilized cardiac fibers

Mitochondrial oxygen consumption was measured using the Oxygraph-2k with an attached fluorometer (OROBOROS Instruments, Innsbruck, Austria). Left ventricular apex tissue was isolated and immediately placed in a 60mm dish containing BIOPS buffer (in mM: CaK_2_<u>EGTA</u> 2.77; K_2_EGTA 7.23; Na_2_ATP 5.77; MgCl_2_*6H_2_O 6.56; taurine 20; Na^2+^2-phosphocreatine 15; <u>imidazole</u> 20; <u>DTT</u> 0.5; MES 50; pH 7.; 4 °C). Cardiac myofibers were gently teased apart using fine-tip tweezers and transferred to a 24 well plate. Fibers were permeabilized using 50 µg/ml saponin in BIOPS for 30 mins. Fibers were washed three times in Miro5 buffer (in mM: 0.5 mM ethylene glycol tetraacetic acid, 3 mM MgCl_2_6H_2_O, 20 mM taurine, 10 mM KH_2_PO_4_, 20 mM N-2-hydroxyethylpiperazine-N-2-ethane <u>sulfonic acid</u>, 110 mM d-sucrose, 0.1% <u>bovine serum</u> <u>albumin</u>and 60 mM <u>lactobionic acid</u>; pH 7.1, 4 degrees) for 10 mins. Fibers were blotted dry, weighed, and immediately added to the Oxygraph chambers. All samples were run in duplicate at 37 °C with a stirring speed of 750 rpm. Hydrogen peroxide emission was measured simultaneously using the fluorimeter with the LED2-Module (525 nM) following the addition of 10 μM Amplex UltraRed, 1 U/mL <u>horseradish peroxidase</u> (HRP) and 5 U/mL superoxide dismutase (SOD). The following SUIT protocol was used to assess mitochondrial bioenergetic function: malate (2 mM), palmitoyl carnitine (5mM) (FAO Leak), ADP (5 mM), Mg^2+^ (5 mM) (FAO OXPHOS), pyruvate (5 mM), glutamate (10mM), ADP (5 mM), Mg^2+^ (5 mM) (FAO + CI OXPHOS), succinate (10 mM) (FAO+CI+CII OXPHOS), ADP (5 mM), Mg^2+^ (5 mM), Oligomycin (2 μg/mL) (FAO+CI+CII Leak). All values were corrected for non-mitochondrial respiration.

### Enzyme activities

Citrate synthase activity was measured in protein lysates, as previously described^75^. Briefly, activity was measured by calculating the rate of absorbance at 412 nM in 50 mM Tris-HCl (pH 8.0), with 0.2 mM DTNB, 0.1 mM acetyl-CoA and 0.25 mM oxaloacetate as the starter chemical using the BioTek Synergy Mx Microplate Reader (BioTek Instruments Inc). Enzyme activity was calculated using the extinction coefficient of 13.6 mM^−1^cm^−1^ for citrate synthase.

### Mitochondrial/nuclear DNA ratio

DNA was isolated from left ventricular tissue as previously described^76^. Briefly, tissue was homogenized in DNA lysis buffer (5 mM EDTA, 0.2% SDS, 200 mM NaCl, 100 mM Tris, pH 8.0) and incubated overnight with proteinase K (ThermoFisher, #25530049). DNA was extracted using phenol/chloroform/isoamyl alcohol (25:24:1; PCIAA) and concentrations were measured using the Nanodrop 2000 (ThermoFisher). qPCR was run using the SsoAdvanced Universal SYBR Green Supermix (Bio-Rad, #1725272) and run on the CFX96 (Bio-Rad) according to manufacturer’s protocol. Mitochondrial-encoded gene (*Nd1* and *16sRNA*) CT values were normalized to nuclear-encoded genes (*Hk2* and *B2m*) using the delta delta CT method^77^. Primer sequences are shown in Table S1.

### RNA Extraction and Quantitative PCR

Flash frozen left ventricle tissue was homogenized in Trizol (ThermoFisher, #15596026) using the MagNA Lyser (Roche, Germany). RNA was isolated according to manufacturers instructions and equalized using Ultrapure H20 (ThermoFisher, #10977015). cDNA was synthesized using the All-In-One 5X RT MasterMix (ABM, #G592). qPCR was performed using the SsoAdvanced Universal SYBR Green Supermix (Bio-Rad, #1725272) and run on the CFX96 (Bio-Rad). Relative transcript expression was calculated using the delta delta CT method^77^. Genes of interest were normalized to the geometric mean of the Ct values of *B2m* and *Eef1e1*. Primer sequences are shown in Table S1.

### Statistical analysis

All statistical tests were computed through their respective Scipy method^62^. Comparison of overall GTEx gene expression was done by Mann-Whitney U test^78^ and OPA1 expression in young and old samples by two-sample T-test^79^. Chi-square test was used to compare pathology occurrence in groups of GTEx histological data^80^.

Unless stated or specified, all data are shown as mean ± SEM. GraphPad Prism software 10.2.3 was used for all statistical analyses of the *in vivo* results. For analyses comparing two groups, an unpaired two-tailed Student’s t-test was used. In all analyses containing two factors (genotype and surgical intervention), data were analyses using a two-way ANOVA with Tukey’s *post-hoc* tests.

## Acknowledgments

This work was funded by a Heart and Stroke Grant-in-Aid to M-E.H. and a New Investigator Award to E.E.M. and Canadian Institute of Health Research (E.E.M. Grant Number 186076, M-E.H. Grant Number 143278). CF-M and SP were supported by Frederick Banting and Charles Best Canada Graduate Scholarships Doctoral Awards. L.K was supported by an Ontario Graduate Scholarship. T.K.T.S. was supported by a Vanier Canada Scholarship. S.M. was supported by an OISB-NRC Graduate Student Scholarship. M.C. was supported by a Faculty of Medicine Summer Studentship Program Award. We thank Dr. Luca Scorrano, Dr. Ruth Slack and Dr. Mireille Khacho for providing us with the OPA1-OE mice; Megan Fortier for technical assistance; and Louise Pelletier Histology Core Facility (RRID: SCR_021737) for technical assistance.

We also acknowledge Conner C. Earl and the Cardiovascular Imaging Research Laboratory Group at Purdue University for their help and training with the 4D ultrasound graphical user interface and strain analysis code.

Schematic figures were created using Biorender.com (publication license: LA277WSPMS).

---

**Supplementary Figure 1.**
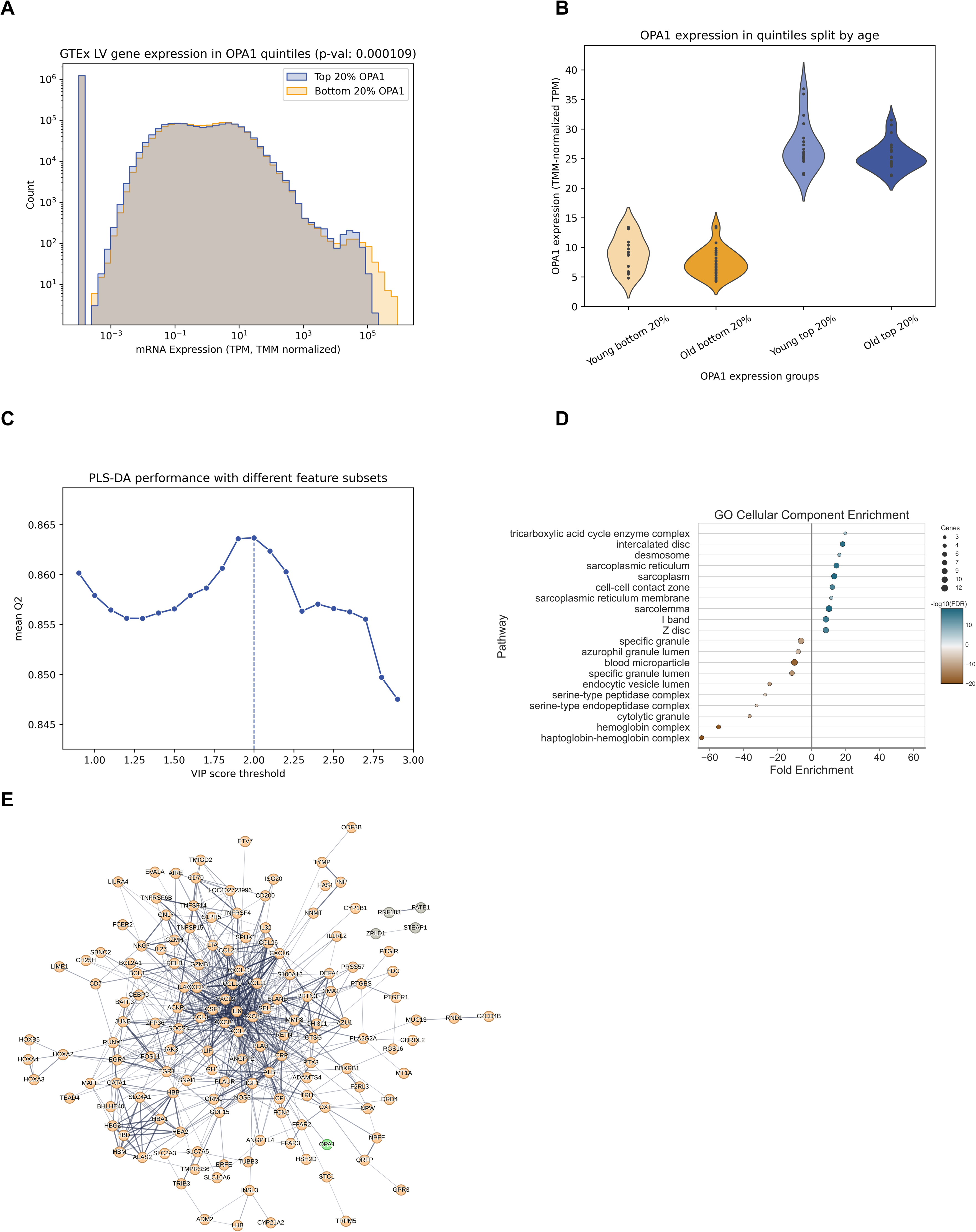
Analysis of OPA1 expression in top and bottom quintiles of GTEx left ventricle samples. (A) Distributions of all gene expression values in top and bottom OPA1 quintiles. **(B)** OPA1 normalized expression values in samples from individuals under 50 (“young”) and 50 or older (“old”) in top and bottom quintiles. **(C)** Performance of PLS-DA classification of top and bottom OPA1 quintile samples by gene expression. Models are trained with decreasing sized gene sets according to their relative importance by VIP score. The best performing model threshold is indicated. **(D)** Enriched GO cellular component terms in positive (blue) and negative (orange) DEGs between top and bottom expressors. **(E)** Protein-protein interaction network of genes negatively associated with OPA1 expression in LV samples.

**Supplementary Figure 2.**
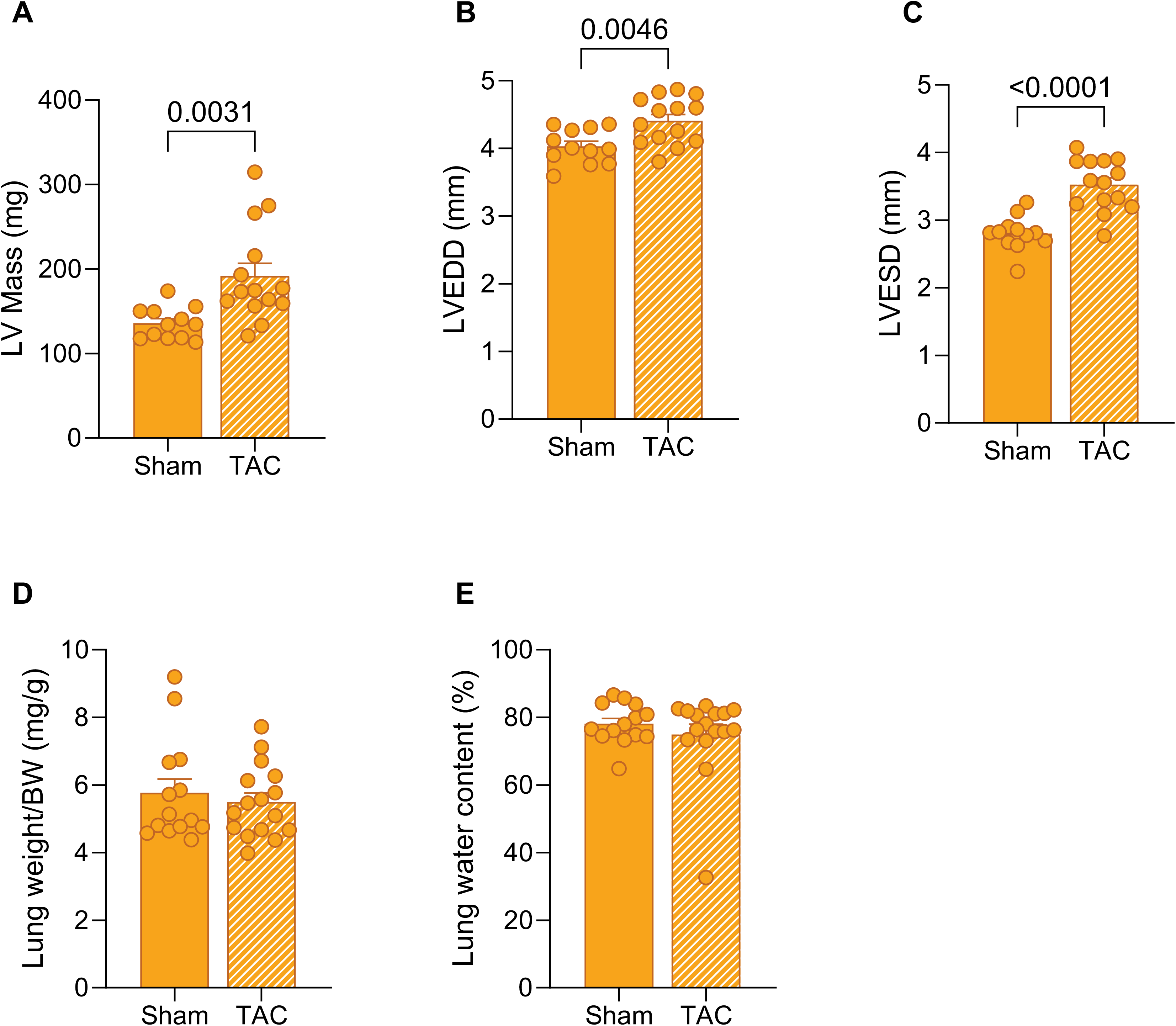
Heart and lung parameters at 12 weeks post TAC. (A-C) Cardiac parameters including left ventricle (LV) mass (A), LV end diastole diameter (B), and LV end systolic diameter (C) were measured using 2-dimensional echocardiography m-mode imaging. **(D)** Lung weight/body weight ratio at 12 weeks post-TAC. **(E)** Lung water content was calculated according to the following equation (wet lung weight – dry lung weight)/wet lung weight × 100%. All data are represented as the mean ± SEM. P values calculated using an unpaired, two-tailed Student’s t-test for (A-E).

**Supplementary Figure 3.**
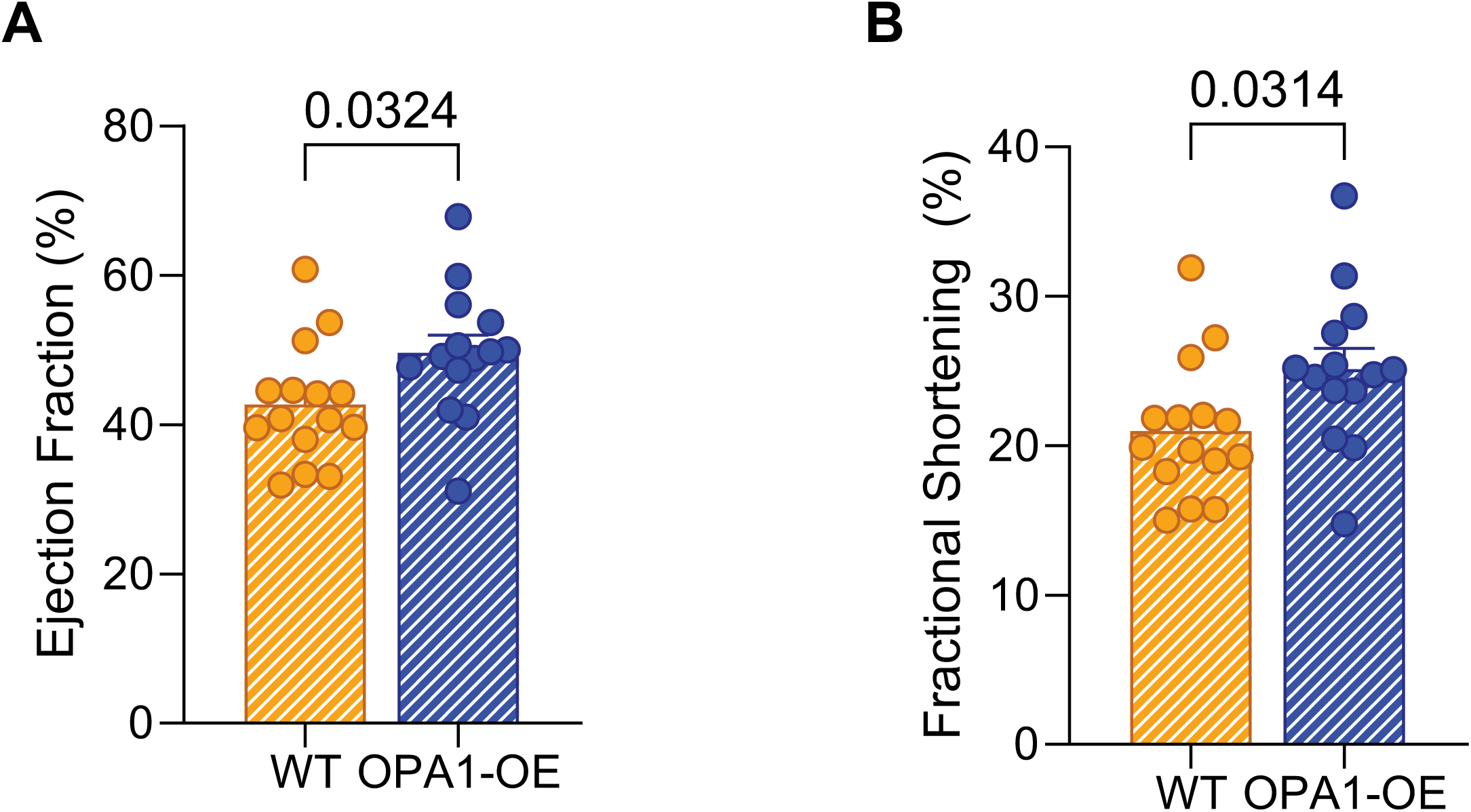
Cardiac function is higher in in OPA1-OE compared to WT TAC mice. (A) Ejection fraction and **(B)** fractional shortening were measured using 2-dimensional echocardiography m-mode imaging. All data are represented as the mean ± SEM. P values calculated using an unpaired, two-tailed Student’s t-test for (A-B).

**Supplemental Table 1.**
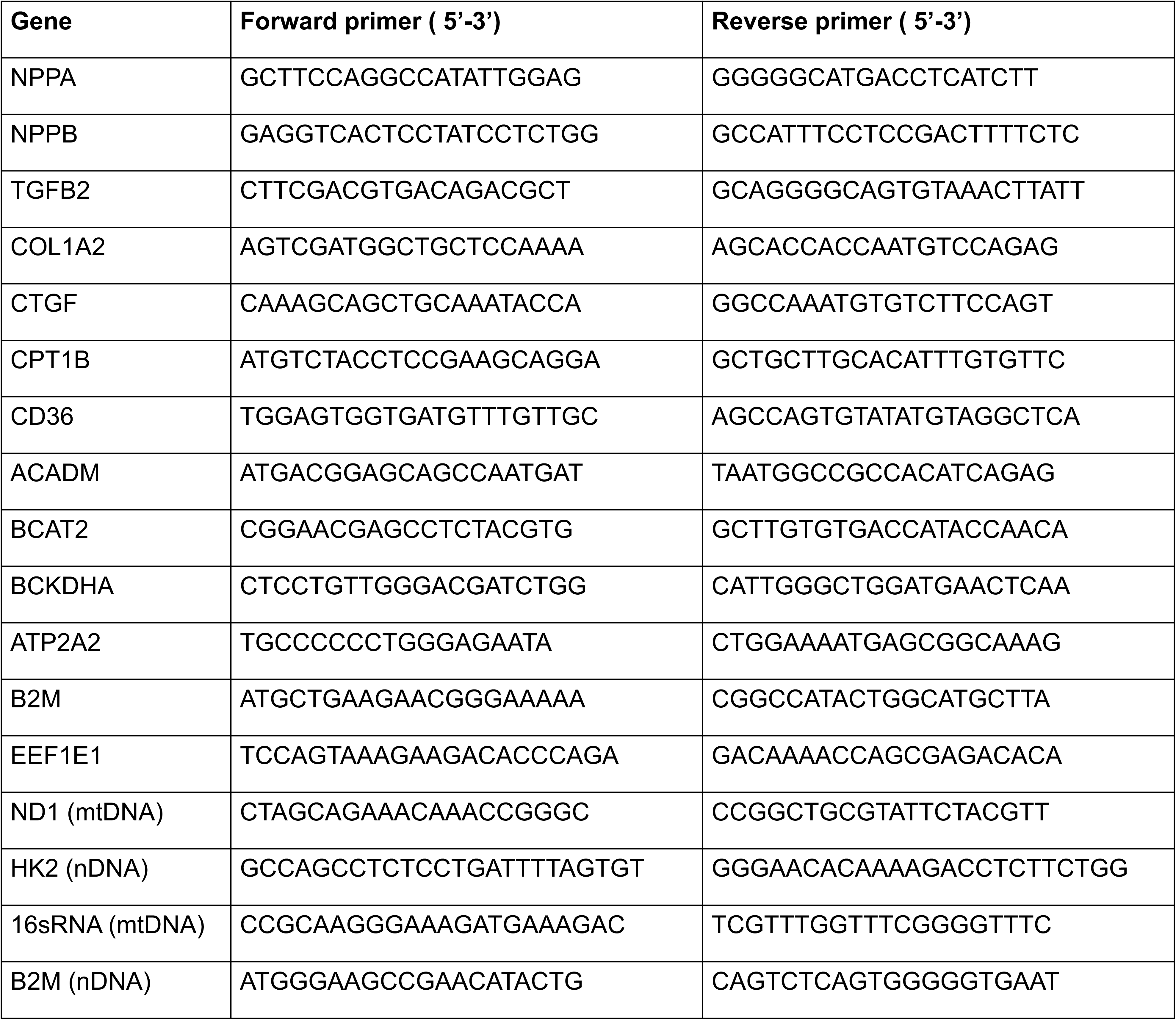
Primer sequences.

